# Mitochondrial superoxide acts in the intestine to extend longevity

**DOI:** 10.1101/2024.09.13.612915

**Authors:** Thomas Liontis, Megan M. Senchuk, Shusen Zhu, Suleima Jacob-Tomas, Ulrich Anglas, Annika Traa, Sonja K. Soo, Jeremy M. Van Raamsdonk

## Abstract

Reactive oxygen species (ROS) are highly reactive oxygen containing molecules that are generated by normal metabolism. While ROS can cause damage to the building blocks that make up cells, these molecules can also act as intracellular signals that promote longevity. The levels of ROS within the cell can be regulated by antioxidant enzymes, such as superoxide dismutase (SOD), which converts superoxide to hydrogen peroxide. Interestingly, our previous work has shown that disruption of the mitochondrial SOD gene *sod-2* results in increased lifespan, indicating that elevating levels of mitochondrial superoxide can promote longevity. To explore the molecular mechanisms involved, we determined the tissues in which disruption of *sod-2* is necessary for lifespan extension and the tissues in which disruption of *sod-2* is sufficient to extend lifespan. We found that tissue-specific restoration of SOD-2 expression in worms lacking SOD-2 could partially revert changes in fertility, embryonic lethality and resistance to stress, but did not inhibit the effects of *sod-2* deletion on lifespan. Knocking down *sod-2* expression using RNA interference specifically in the intestine, but not other tissues, was sufficient to extend longevity. Intestine-specific knockdown of *sod-2* also increased resistance to heat stress and while decreasing resistance to oxidative stress. Combined, these results indicate that disruption of *sod-2* in neurons, intestine, germline, or muscle is not required for lifespan extension, but that decreasing *sod-2* expression in just the intestine extends lifespan. This work defines the conditions required for elevated mitochondrial superoxide to increase longevity.

## Introduction

Superoxide is the main form of reactive oxygen species (ROS) generated by mitochondria as a by-product of cellular respiration. Based on the observation that oxidative damage increases with age, the Free Radical Theory of Aging (FRTA) proposed that oxidative damage caused by ROS contributes to aging [1]. At the same time, ROS can act as intracellular signaling molecules to activate pathways that protect against stress and promote longevity. To control the levels of ROS in a cell, organisms have multiple antioxidant enzymes. Superoxide dismutase (SOD) is the only eukaryotic enzyme that is able to detoxify superoxide. In *C. elegans*, *sod-1, sod-2*, and *sod-4* (*SOD1*, *SOD2*, and *SOD3* in humans) encode the primary cytoplasmic, mitochondrial, and extracellular SODs, respectively. *C. elegans* also expresses two additional *sod* genes: *sod-3* and *sod-5,* which are expressed in the mitochondria and cytoplasm at low levels.

We and others have previously examined the effect of disrupting individual or combinations of *sod* genes on lifespan. While the FRTA predicts that increasing ROS levels would decrease lifespan, surprisingly, deletion of individual *sod* genes or all five *sod* genes together have little or no detrimental effect on lifespan despite increasing sensitivity to oxidative stress [2–5]. In fact, we found that deletion of the mitochondrial *sod* gene *sod-2* can significantly increase lifespan [4]. Similarly, directly increasing mitochondrial superoxide levels through treatment with low concentrations of the superoxide-generating compounds paraquat or juglone can also extend longevity [6, 7]. In addition, increased ROS levels have been shown to be required for the longevity of other long-lived mutants including *daf-2* insulin/IGF-1 receptor mutants [8], *glp-1* germline ablation mutants [9] and multiple mutants with mild impairment of mitochondrial function [6, 10].

There is now an increasing amount of evidence that points to a beneficial role of ROS for prolonging life that is conserved across species. Indeed, interventions or mutations that increase ROS have also been shown to extend longevity in yeast [11, 12], flies [13, 14], and mice [15, 16]. While the effect of ROS on longevity in humans has not been examined, elevation in ROS has been shown to contribute to the healthspan-promoting effects of exercise [17]. Furthermore, in a meta-analysis of studies in which humans were treated with various antioxidants, there was no consistent beneficial effect of antioxidants on human longevity [18].

In order to fully understand the mechanisms by which mild elevation of mitochondrial superoxide extends longevity, it is first necessary to define the conditions under which superoxide promotes longevity. Performing mechanistic studies in whole worms may mask important molecular events that are occurring in specific tissues, thereby complicating the understanding of the physiological relationship between mitochondrial superoxide and lifespan. Accordingly, in this work we identify the tissues in which mitochondrial superoxide is acting to increase lifespan.

Previous experiments have found that genetic modifications in a single tissue can increase the lifespan of the whole organism [19–22], indicating the occurrence of cell non-autonomous effects on lifespan. In long-lived *daf-2* mutants, expressing wild-type *daf-2* in neurons completely suppresses their long life, indicating that disruption of *daf-2* in neurons is necessary for their lifespan extension [23]. Tissue-specific overexpression the FOXO transcription factor DAF-16 in the intestine increases the lifespan of *daf-2;daf-16* double mutants, indicating that expression of *daf-16* in the intestine is sufficient to enhance longevity [24]. Similarly, disrupting subunits of the mitochondrial electron transport chain in either intestine or neurons is sufficient to extend the lifespan of the entire organism [20].

In this work we determine the tissues in which *sod-2* disruption is necessary or sufficient to extend longevity, alter stress resistance and affect physiologic rates. Our results suggest that there is no single tissue in which *sod-2* deletion is required to increase lifespan, but that knockdown of *sod-2* specifically in the intestine is sufficient to enhance longevity. These results highlight the importance of the intestine in determining longevity in a cell non-autonomous manner.

## Materials and Methods

### Strains

**N2 (CGC)** wild-type

**MQ130** *clk-1(qm30)*

**WM27** *rde-1(ne219)*

**NL2098** *rrf-1(pk1417)*

**NL3321** *sid-1(pk3321)*

**JVR501** *clk-1(qm30); rde-1(ne219)*

**JVR503** *clk-1(qm30); sid-1(pk3321)*

**JVR561** *clk-1(qm30); rrf-1(pk1417)*

**NR222** *rde-1(ne219)*; *kzIs9[pKK1260(lin-26p::nls::GFP) + pKK1253(lin-26p::rde-1) + pRF6(rol-6(su1006)]*

**NR350** *rde-1(ne219); kzIs20[pDM#715(hlh-1p::rde-1) + pTG95(sur-5p::nls::GFP)]*

**VP303** *rde-1(ne219); kbIs7[nhx-2p::rde-1 + rol-6(su1006)]*

**TU3401** *sid-1(pk3321); uIs69[pCFJ90(myo-2p::mCherry) + unc-119p::sid-1] V*

**MQ1503** *clk-1(qm30); sod-2(ok1030)*

**MQ1451** *sod-2(ok1030)*

**JVR348** *clk-1(qm30); sod-2(ok1030); jerIs003[sod-2p::SOD-2 genomic::eGFP::let-858utr + unc-119(+)]*

**JVR468** *clk-1(qm30); sod-2(ok1030); jerIs006[pie-1p::SOD-2 genomic::eGFP::let-858utr + unc-119(+)]*

**JVR469** *clk-1(qm30); sod-2(ok1030); jerIs007[myo-3p::SOD-2 genomic::eGFP::let-858utr + unc-119(+)]*

**JVR471** *clk-1(qm30); sod-2(ok1030); jerIs011[elt-2p::SOD-2 genomic::eGFP::let-858utr + unc-119(+)]*

**JVR472** *clk-1(qm30); sod-2(ok1030); jerIs012[unc-119p::SOD-2 genomic::eGFP::let-858utr + unc-119(+)]*

**JVR565** *clk-1(qm30); sid-1(pk3321); uIs69[pCFJ90(myo-2p::mCherry) + unc-119p::sid-1] V*

**JVR566** *clk-1(qm30); rde-1(ne219); kbIs7[nhx-2p::rde-1 + rol-6(su1006)]*

**JVR567** *clk-1(qm30); rde-1(ne219); kzIs20[pDM#715(hlh-1p::rde-1) + pTG95(sur-5p::nls::GFP)]*

**JVR568** *clk-1(qm30); rde-1(ne219)*; *kzIs9[pKK1260(lin-26p::nls::GFP) + pKK1253(lin-26p::rde-1) + pRF6(rol-6(su1006)]*

### Generation of tissue-specific *sod-2* expression strains

The *sod-2* gene was cloned from genomic DNA into pDONR221 using the following attB site containing primers.

Forward: 5’ GGGGACAAGTTTGTACAAAAAAGCAGGCTtt CTT CAA AAC ACC GTT CGC TGT GTC TC Reverse 5’ GGGGACCACTTTGTACAAGAAAGCTGGGTt TTG CTG TGC CTT TGC AAA ACG C

Start and stop codons were contributed in-frame by 5’ and 3’ elements. LR multisite Gateway cloning was used to generate tissue-specific *sod-2* constructs using promoter constructs obtained through the *C. elegans* Promoterome project (*sod-2, myo-3, unc-119, elt-2*) or Addgene (*pie-1*, pCM1.58). 3’ eGFP::let-858utr (bsem1157) and destination vector (pCFJ150) were provided courtesy of the Mango and Jorgensen labs, as were additional MosSCI (Mos1-mediated single copy insertion) reagents. *sod-2* constructs were injected at 25 ng/µl and integrated by MosSCI at ttTi5605 (II:0.77) using standard methods (Frøkjær-Jensen et al, 2014). Strains were outcrossed prior to characterization.

### RNA isolation and quantitative RT-PCR

To isolate RNA, pre-fertile young adult worms from a limited lay were collected in M9 buffer, washed three times and then frozen in TRIZOL. mRNA was isolated as previously described [25] and then converted to cDNA using a High-Capacity cDNA Reverse Transcription Kit (Life Technologies). A FastStart Universal SYBR Green kit (Roche) was used to perform Quantitative RT-PCR in an AP Biosystems RT-PCR machine. The following primers were used:

*sod-2* Forward: GCTATTTGGAAGATCGCC

*sod-2* Reverse: TAGTTACAAAACAATGGGAGTC

*egfp* Forward: GTTCCATGGCCAACACTTGT

*egfp* Reverse: GACTTCAGCACGTGTCTTGTAG

### Fertility and embryonic lethality

Fertility was measured by quantifying the self-brood size of 8-15 animals per genotype or condition. Individual animals at the L4 stage were moved to their own NGM plate. Once egg laying had begun, these animals were transferred to new plates daily or every two days until egg laying ceased. The brood size was counted as the number of live animals that hatched from each worm. Embryonic lethality was measured by performing a 3-hour limited lay. After 2 days the number of unhatched eggs and live worms was counted. Embryonic lethality was calculated as the number of unhatched eggs divided by the total number of eggs laid.

### Post-embryonic development time

Post-embryonic development time was measured by transferring at least 200 eggs to a fresh NGM plate seeded with OP50 bacteria. After three hours, approximately 20-25 L1 worms that had hatched during the three-hour window were transferred to a new plate. After 20 hours, these worms were checked every 4 hours. Worms that reached adulthood were counted and removed. Three biological replicates were performed.

### Movement – thrashing rate

Movement was measured as the number of thrashes per second in M9 buffer. Worms were synchronized by bleaching and allowed to grow to adulthood in NGM plates seeded with OP50 bacteria. At adulthood, M9 buffer was added to the plate. After 1 minute, a video of the worms thrashing was captured using WormLab (Microbrightfield). The 1-minute-long videos were analyzed using the wrMTrck plugin for Fiji. Three biological replicates were performed.

### RNA interference

In order to induce RNA interference (RNAi), worms were transferred to RNAi plates, which consist of NGM (nematode growth medium) supplemented with the antibiotic carbenicillin (25 μg/ml) and IPTG (1mM). Worms were transferred to RNAi plates at the L4 stage of the parental generation. Once egg laying began, gravid adults were transferred to a new RNAi plate for a 24-hour limited lay. These progeny were used for the lifespan assay.

### Lifespan assay

Lifespan assays were conducted on NGM plates supplemented with 5-fluorodeoxyuridine (FUdR) [26]. Although FUdR can affect the lifespan of some strains [27, 28], it does not affect the lifespan of wild-type worms and at low concentrations of 25 µM or less it has minimal effects on longevity. Accordingly, lifespan assays were performed at 25 µM FUdR to minimize effects on lifespan while maximizing inhibition of progeny development. For lifespan assays, 3 independent biological replicates, each containing at least 50 worms, were performed. Deaths were scored every 2-3 days. Worms were considered dead when they stopped exhibiting spontaneous movement and failed to move in response to 1) a gentle touch to their tail, 2) a gentle touch to their head, and 3) a gentle lifting of their head. If an animal died by non-natural causes, i.e. from internal hatching or from their intestine bursting or leaking, it was censored.

### Chronic oxidative stress assay

Resistance to chronic oxidative stress was assessed by exposing worms to 4 mM paraquat and checking survival daily. These plates contained 100 µM FUdR as paraquat treatment induces internal hatching. Three biological replicates with at least 20 worms per replicate were performed.

### Heat stress assay

Resistance to heat stress was measured by exposing worms to 37°C heat stress for 3 hours followed by a 24-hour recovery period. Four biological replicates with at least 20 worms per replicate were performed.

### Bacterial pathogen stress assay

Resistance to bacterial pathogens was assessed by examining survival after exposure to *P. aeruginosa* strain PA14 in a slow kill assay [29, 30]. Briefly, young adult worms were grown on plates containing 100 mg/L FUdR until day 3 of adulthood. At this point, worms were transferred to plates containing 20 mg/L FUdR, which were seeded with *P. aeruginosa* strain PA14, and survival was monitored daily until death at 20°C. Three biological replicates with 20-30 worms per replicate were performed.

### Statistical analysis

Statistical analyses were performed using GraphPad Prism version 9. For column graphs, we used a one-way ANOVA with Dunnett’s multiple comparison test. For assays with multiple time points, we used a two-way ANOVA with Bonferroni’s multiple comparisons test. For survival assays, we used a log-rank test. Error bars indicate standard error of the mean (SEM). *p<0.05, **p<0.01, ***p<0.001, ****p<0.0001.

## Results

### Expression of SOD-2 in specific tissues is sufficient to rescue deficits in fertility and embryonic lethality caused by global disruption of *sod-2*

Our previous work has shown that disruption of the mitochondrial *sod* gene, *sod-2*, results in the alteration of several physiologic rates, modulation of stress resistance and extension of lifespan, and that these phenotypic differences are enhanced in a *clk-1* ROS-sensitive background [4, 31]. To better understand the underlying mechanisms involved, we sought to identify the tissues in which disruption of *sod-2* is necessary for the observed alterations in phenotype. To do this, we generated novel transgenic worm strains that overexpress SOD-2 linked to enhanced green fluorescent protein (eGFP) either ubiquitously or in specific tissues using tissue-specific promoters. For ubiquitous expression, we used the endogenous *sod-2* promoter, while expression in the germline, muscle, intestine and neurons was achieved using the *pie-1, myo-3, elt-2* and *unc-119* promoters, respectively. The *sod-2* overexpression strains were generated using Mos1-mediated single copy insertion (MosSCI) in order to achieve expression levels comparable to endogenous [32]. The tissue-specific *sod-2* expression strains were then crossed to *clk-1;sod-2* worms.

As a first step, we examined where in the worm *sod-2* was being expressed by visualizing eGFP expression. In the ubiquitous strain (*clk-1;sod-2;sod-2p::sod-2::egfp*), SOD-2::eGFP could be seen throughout the worm (**Figure 1A**). In the germline-specific strain (*clk-1;sod-2;pie-1p::sod-2::egfp*), SOD-2::eGFP expression was limited to the germline (**Figure 1A**). In the muscle-specific strain (*clk-1;sod-2;myo-3p::sod-2::egfp*), SOD-2::eGFP expression was limited to the body wall muscle (**Figure 1A**). In the intestine-specific strain (*clk-1;sod-2;elt-2p::sod-2::egfp*), SOD-2::eGFP expression was limited to the intestine (**Figure 1A**). In the neuron-specific strain (*clk-1;sod-2;unc-119p::sod-2::egfp*), SOD-2::eGFP expression was limited to neurons (**Figure 1A**).

**Figure 1.**
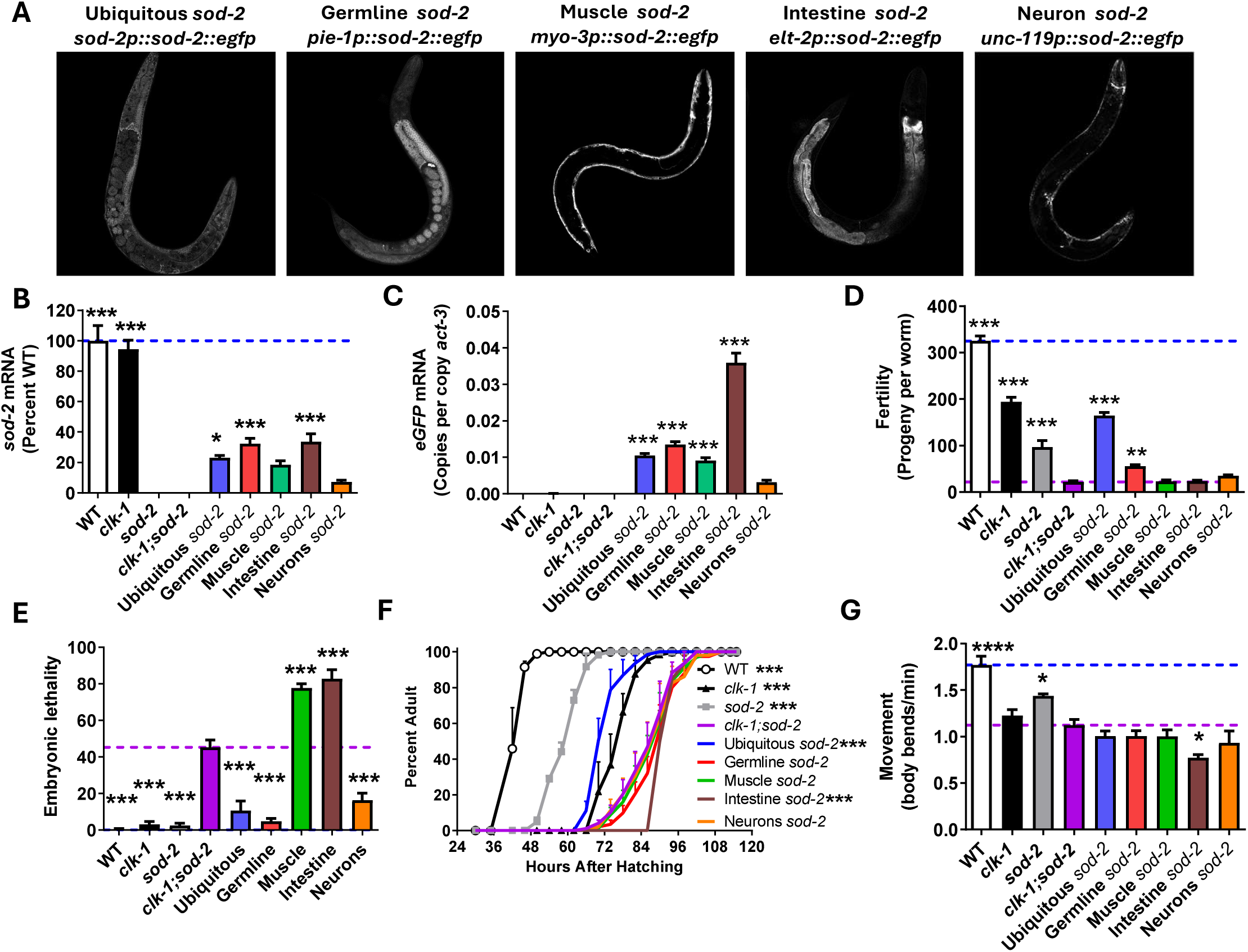
Tissue-specific overexpression of *sod-2* rescues deficits in movement, fertility and embryonic lethality in *clk-1;sod-2* mutants. Transgenic *C. elegans* strains overexpressing *sod-2::egfp* under tissue-specific promoters exhibit fluorescence from eGFP in the appropriate tissue indicating that the expression of *sod-2::egfp* is being limited to the targeted tissue (**A**). Measurement of whole worm *sod-2* mRNA levels by quantitative RT-PCR (qPCR) demonstrates that *sod-2* is being expressed in the tissue-specific rescue strains but at a lower level than in wild-type worms (**B**). Similarly, measurement of *egfp* mRNA levels by qPCR shows that the *sod-2::egfp* transgene is being expressed in the tissue-specific rescue strains (**C**). Ubiquitous or germline-specific expression of *sod-2::egfp* increases brood size in *clk-1;sod-2* worms (**D**). Embryonic lethality in *clk-1;sod-2* worms is reduced by ubiquitous, germline or neuronal expression of *sod-2::egfp* but exacerbated by muscle- or intestine-specific expression of *sod-2::egfp* (**E**). Only ubiquitous expression of *sod-2::egfp* is sufficient to rescue slow development time of *clk-1;sod-2* worms (**F**). Intestine-specific expression of *sod-2::egfp* decreases thrashing rate in *clk-1;sod-2* worms (**G**). Statistical significance indicates differences from *clk-1;sod-2* worms. Statistical significance was assessed using a one-way ANOVA with Dunnett’s multiple comparison test in panels B-E,G and a repeated measures ANOVA with Bonferroni’s multiple comparisons test in panel F. Ubiquitous *sod-2* = *clk-1;sod-2;sod-2p::sod-2::egfp*, Germline *sod-2* = *clk-1;sod-2;pie-1p::sod-2::egfp,* Muscle *sod-2* = *clk-1;sod-2;myo-3p::sod-2::egfp,* Intestine *sod-2* = *clk-1;sod-2;let-2p::sod-2::egfp,* Neurons *sod-2* = *clk-1;sod-2;unc-119p::sod-2::egfp.* Blue dotted line indicates wild-type level, purple dotted line indicates *clk-1;sod-2* level. Error bars indicate standard error of the mean. *p<0.05,**p<0.01, ***p<0.001,****p<0.0001.

Next, we examined the expression levels of *sod-2* and *egfp* using quantitative RT-PCR. As expected, *sod-2* expression was absent in *sod-2* and *clk-1;sod-2* worms (**Figure 1B**). The expression of *sod-2* was detected in all of the tissue-specific rescue strains, but at a level that is lower than in wild-type worms (**Figure 1B**). The level of *sod-2* expression in the tissue-specific strains was not significantly different from the ubiquitous rescue strain, except in the case of the neuron-specific rescue strain. *Egfp* expression was detected in all of the *sod-2::egfp* expression strains (**Figure 1C**).

Having confirmed that the rescue strains were expressing *sod-2::egfp* in the correct tissues, we sought to determine whether this would be sufficient to rescue deficits in physiologic rates in *clk-1;sod-2* worms. *clk-1;sod-2* mutants have a markedly decreased brood size, which was rescued in the ubiquitous rescue strain (**Figure 1D**). Among the tissue-specific rescue strains, only expression of *sod-2::egfp* in the germline was able to increase the brood size in *clk-1;sod-2* mutants (**Figure 1D**). In addition to having a decreased brood size, *clk-1;sod-2* mutants have markedly increased embryonic lethality (**Figure 1E**). This increase in embryonic lethality was prevented by expression of *sod-2::egfp* ubiquitously, or specifically in germline or neurons (**Figure 1E**). In contrast, the muscle-specific and intestine-specific rescues strains exhibited a significant increase in embryonic lethality (**Figure 1E**).

Next, we examined postembryonic development time. While ubiquitous expression of *sod-2::egfp* was able to completely rescue the slow development time of *clk-1;sod-2* worms, all of the tissue-specific rescue strains exhibited a development time that was equivalent to or slower than *clk-1;sod-2* worms (**Figure 1F**). Finally, the thrashing rate of *clk-1;sod-2* worms was not increased by *sod-2* expression either ubiquitously or in specific tissues, but was decreased under intestine-specific SOD-2 rescue (**Figure 1G**).

Overall, germline-specific and neuronal-specific rescues of SOD-2 were beneficial in preventing developmental viability deficits of *clk-1; sod-2* worms, whereas intestine-specific and muscle-specific rescues of SOD-2 exacerbated them. Intestinal-specific rescue of SOD-2 also impaired movement.

### Expression of SOD-2 in specific tissues modulates altered stress resistance caused by global disruption of *sod-2*

To determine the tissues in which disruption of *sod-2* is necessary for altering resistance to stress, we characterized resistance to bacterial pathogen stress, heat stress and oxidative stress in the tissue-specific rescue strains. We found that *clk-1;sod-2* mutants have increased resistance to *P. aeruginosa* strain PA14 in a slow kill assay compared to wild-type or *clk-1* worms (**Figure 2A**). This enhanced resistance to bacterial pathogens was prevented by ubiquitous expression of *sod-2* and significantly decreased by expression of *sod-2* in the germline or neurons (**Figure 2A**). *clk-1;sod-2* worms also exhibit increased resistance to heat stress compared to wild-type and *clk-1* worms (**Figure 2B**). Re-expression of *sod-2* in either the body wall muscle or neurons significantly decreased survival under heat stress (**Figure 2B**). Finally, we found that ubiquitous expression of *sod-2* decreased resistance to chronic oxidative stress in *clk-1;sod-2* worms to wild-type level (**Figure 2C**). In contrast, re-expression of *sod-2* in the muscle, intestine or neurons of *clk-1;sod-2* mutants all resulted in increased survival under oxidative stress (**Figure 2C**).

**Figure 2.**
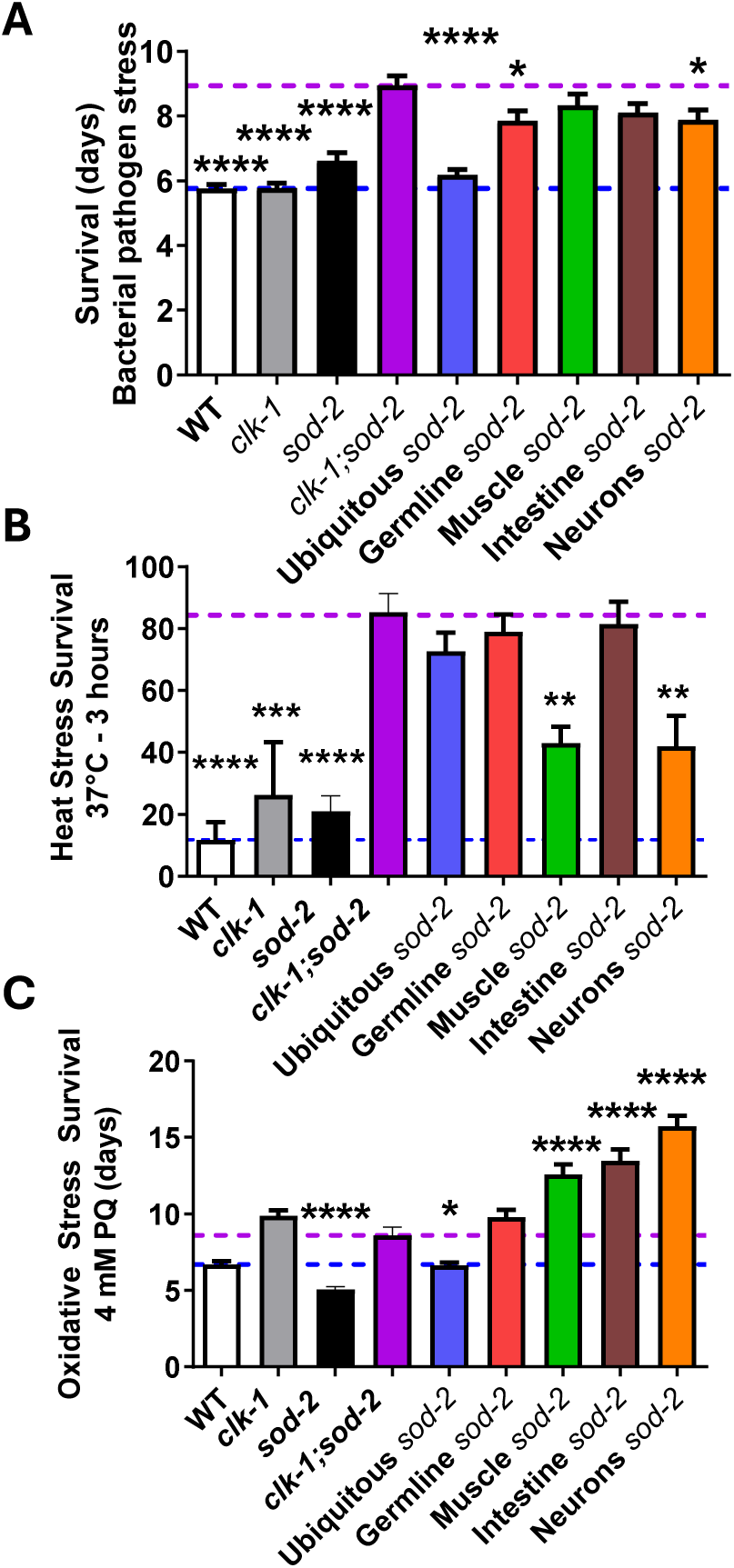
Tissue-specific overexpression of *sod-2* can decrease stress resistance in *clk-1;sod-2* worms. **(A)** Ubiquitous expression of *sod-2::egfp* decreased *clk-1;sod-2* resistance to bacterial pathogens *(P. aeruginosa* strain PA14). A small but significant decrease was also observed for expression of *sod-2::egfp* in germline or neurons. **(B)** Muscle- or neuron-specific expression of *sod-2::egfp* decreases *clk-1;sod-2* worms’ enhanced resistance to heat stress (37°C). **(C)** Ubiquitous expression of *sod-2::egfp* decreased resistance to chronic oxidative stress (4 mM paraquat) in *clk-1;sod-2* worms, while expression in the muscle, intestine or neurons increased survival. Statistical significance indicates differences from *clk-1;sod-2* worms. Statistical significance was assessed using a one-way ANOVA with Dunnett’s multiple comparison test in panels A-C. Ubiquitous *sod-2* = *clk-1;sod-2;sod-2p::sod-2::egfp*, Germline *sod-2* = *clk-1;sod-2;pie-1p::sod-2::egfp,* Muscle *sod-2* = *clk-1;sod-2;myo-3p::sod-2::egfp,* Intestine *sod-2* = *clk-1;sod-2;let-2p::sod-2::egfp,* Neurons *sod-2* = *clk-1;sod-2;unc-119p::sod-2::egfp.* Blue dotted line indicates wild-type level, purple dotted line indicates *clk-1;sod-2* level. Error bars indicate standard error of the mean. *p<0.05,**p<0.01, ***p<0.001,***p<0.0001.

### Expression of SOD-2 in specific tissues is not sufficient to limit lifespan extension caused by global disruption of *sod-2*

Next, we sought to identify tissues in which disruption of *sod-2* is necessary for lifespan extension. First, we confirmed that *sod-2* mutants live significantly longer than wild-type worms and exhibit a markedly increased lifespan in a *clk-1* ROS-sensitive background (**Figure 3A**). We also confirmed that ubiquitous expression of *sod-2::egfp* decreases *clk-1;sod-2* lifespan such that the lifespan of the ubiquitous rescue strain is identical to *clk-1* (**Figure 3B**). In contrast, we found that expression of *sod-2::egfp* in germline (**Figure 3C**), muscle (**Figure 3D**), intestine (**Figure 3E**) or neurons (**Figure 3F**) did not decrease *clk-1;sod-2* lifespan. Intriguingly, neuron-specific, muscle-specific, or germline-specific rescue of SOD-2 instead resulted in a small increase in lifespan. These results suggest that there is no specific tissue where *sod-2* needs to be absent to extend lifespan, but rather disruption of *sod-2* from multiple tissues may contribute to lifespan extension. Alternatively, it is possible that disruption of *sod-2* in a tissue that we did not examine is necessary for lifespan extension in *clk-1;sod-2* mutants.

**Figure 3.**
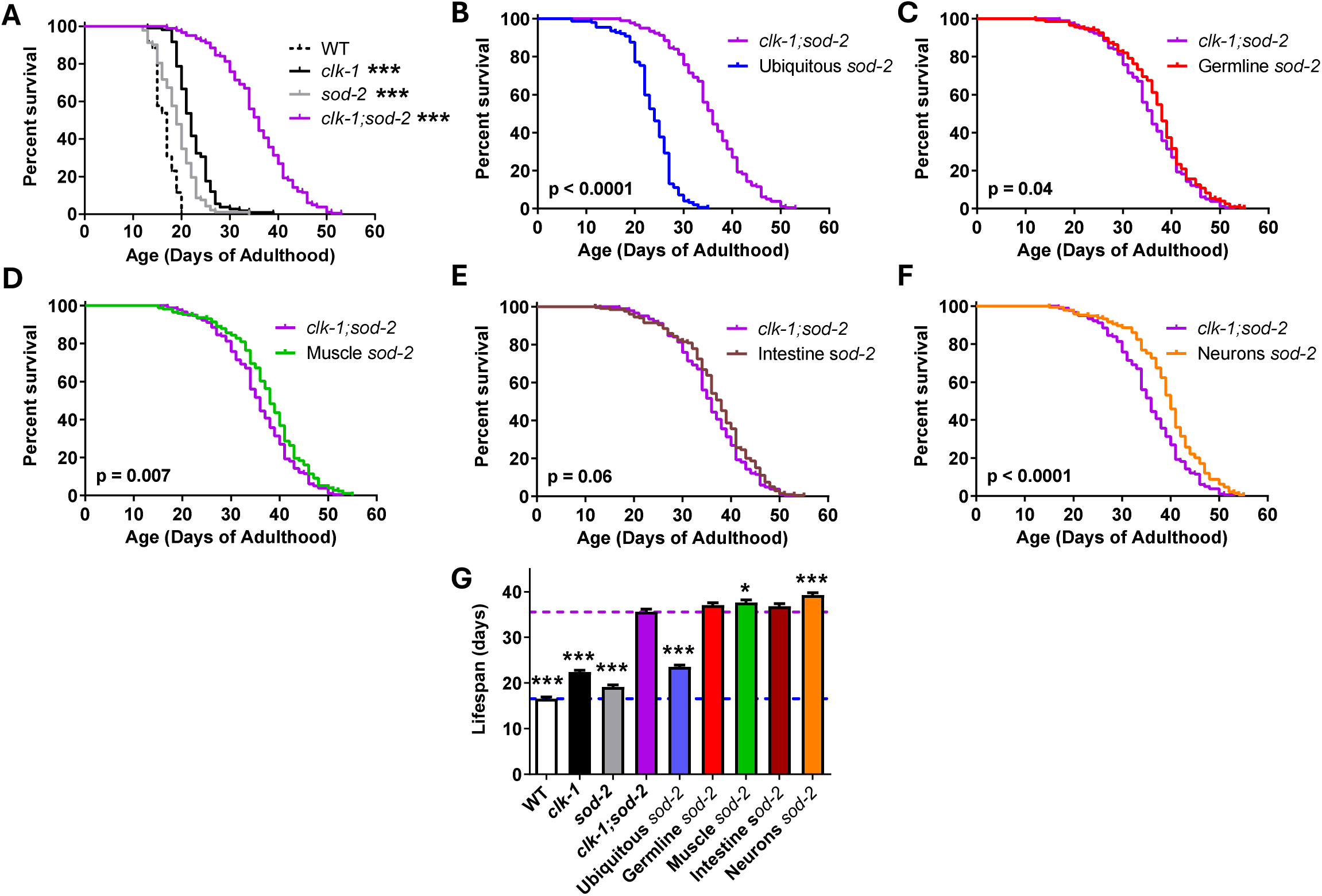
Tissue-specific overexpression of *sod-2* is not sufficient to reduce *clk-1;sod-2* lifespan. Deletion of *sod-2* increases lifespan in wild-type and *clk-1* backgrounds. The magnitude of lifespan extension is much greater in *clk-1* worms (**A**). Ubiquitous expression of *sod-2::egfp* decreases *clk-1;sod-2* lifespan to *clk-1* lifespan (**B**). In contrast tissue-specific expression of *sod-2::egfp* in germline (**C**), muscle (**D**), intestine (**E**), or neurons (**F**) does not decrease *clk-1;sod-2* lifespan. Bar graph showing lifespan of all strains (**G**). Statistical significance indicates differences from *clk-1;sod-2* worms. Statistical significance was assessed using a log-rank test in panels A-F and a one-way ANOVA with Dunnett’s multiple comparison test in panel G. Ubiquitous *sod-2* = *clk-1;sod-2;sod-2p::sod-2::egfp*, Germline *sod-2* = *clk-1;sod-2;pie-1p::sod-2::egfp,* Muscle *sod-2* = *clk-1;sod-2;myo-3p::sod-2::egfp,* Intestine *sod-2* = *clk-1;sod-2;let-2p::sod-2::egfp,* Neurons *sod-2* = *clk-1;sod-2;unc-119p::sod-2::egfp.* Blue dotted line indicates wild-type level, purple dotted line indicates *clk-1;sod-2* level. Error bars indicate standard error of the mean. *p<0.05, **p<0.01, ***p<0.001.

### Disruption of *sod-2* in the intestine is sufficient to extend longevity

Of the tissues that we examined, there was no tissue in which the disruption of *sod-2* is necessary for lifespan extension. Next, we determined whether disruption of *sod-2* in specific tissues is sufficient to extend lifespan. For this purpose, we used tissue-specific RNAi strains, which had been generated and characterized previously [20, 33–35]. In these strains, a required component of the RNAi machinery is expressed using a tissue-specific promoter in a mutant that is globally deficient for that specific component. Specifically, to achieve RNAi knockdown specifically in germline, muscle, intestine and hypodermis, we expressed the Argonaute protein RDE-1, which is required for RNAi, in *rde-1* deficient worms using tissue-specific promoters (*myo-3p* for muscle, *nhx-2p* for intestine, *lin-26p* for hypodermis), using strains that were previously validated [20, 33, 34]. For tissue-specific expression in neurons, we expressed the dsRNA import channel SID-1 under the pan-neuronal *unc-119* promoter in a *sid-1* mutant background [35]. For germline-specific RNAi, we used the *rrf-1* mutation which selectively decreases somatic RNAi [36–38]). All of these tissue-specific RNAi strains were crossed into a *clk-1* mutant background to enhance our chances of seeing an effect and then treated with *sod-2* RNAi.

We found that *sod-2* RNAi significantly increases lifespan in *clk-1* worms (**Figure 4A**), but not wild-type (**Figure 4B**). *sod-2* RNAi has no effect on *clk-1;rde-1* mutants indicating that the *rde-1* mutation effectively inhibits RNAi (**Figure 4C**). Intestine-specific knockdown of *sod-2* is sufficient to increase *clk-1* lifespan (**Figure 4D**), while neuron-specific or germline-enriched knockdown of *sod-2* had no effect on *clk-1* lifespan (**Figure 4E,F**). Knocking down *sod-2* expression in the muscle or hypodermis resulted in a small but significant decrease in *clk-1* lifespan (**Figure 4G,H**). Combined, this indicates that disruption of *sod-2* specifically in the intestine is sufficient to extend lifespan.

**Figure 4.**
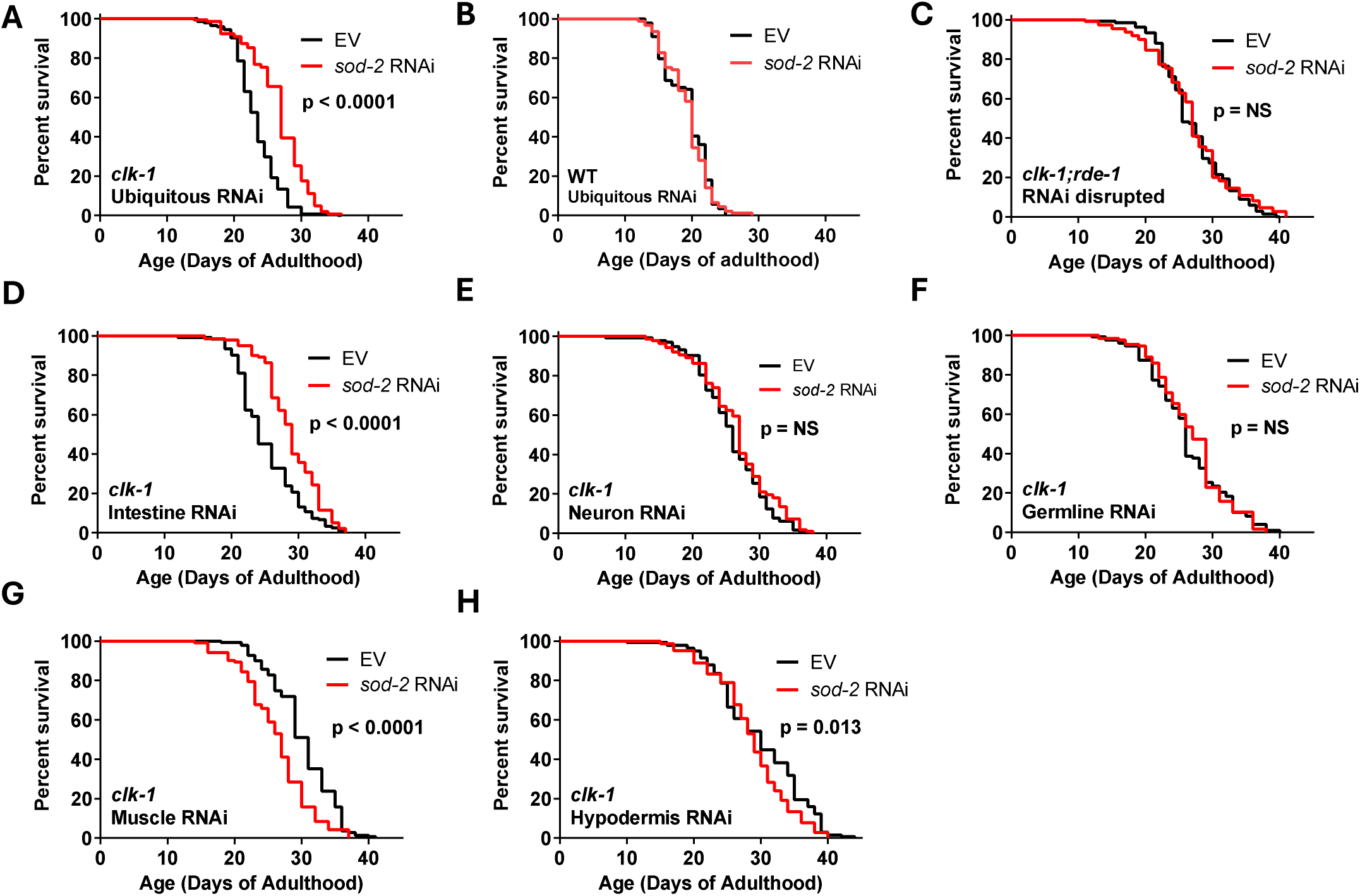
Mild elevation of mitochondrial superoxide in the intestine is sufficient to increase lifespan. To identify the tissues in which disruption of *sod-2* is sufficient to extend lifespan, we crossed tissue-specific RNAi strains to *clk-1* mutants and then knocked down *sod-2* expression using RNAi. Global knockdown of *sod-2* expression increases lifespan in *clk-1* worms (**A**) but not wild-type (**B**). Treating *clk-1;rde-1* worms with *sod-2* RNAi has no effect on longevity indicating that the *rde-1* mutation inhibits RNAi (**C**). Knocking down *sod-2* expression in the intestine is sufficient to increase *clk-1* lifespan (**D**). Neuron-specific or germline-specific knockdown of *sod-2* has no effect on *clk-1* lifespan (**E,F**) while muscle-specific or hypodermis-specific knockdown of *sod-2* results in a small decrease in *clk-1* lifespan (**G,H**). Three biological replicates were performed. Statistical significance was assessed using the log rank test. Germline RNAi = *clk-1;rde-1;mkcIs13[sun-1p::rde-1],* Muscle RNAi = *clk-1;rde-1;kzIs20[hlh-1p::rde-1 + sur-5p::nls::GFP],* Intestine RNAi = *clk-1;rde-1;kbIs7[nhx-2p::rde-1 + rol-6],* Hypodermis RNAi = *clk-1;rde-1;kzIs9[lin-26p::rde-1 + lin-26p::nls::GFP + rol-6],* Neuron RNAi = *clk-1;sid-1;uIs69[unc-119::sid-1 + myo-2p::mCherry]*.

### Tissue-specific knockdown of *sod-2* modulates resistance to stress

We have previously shown that global disruption of *sod-2* increases resistance to bacterial pathogens and heat stress, while decreasing resistance to oxidative stress [4, 39]. Accordingly, we next examined the extent to which knockdown of *sod-2* in the intestine is sufficient to alter resilience in a *clk-1* ROS-sensitive background. We used three different stress paradigms: exposure to bacterial pathogens (*P. aeruginosa* strain PA14, slow kill assay), exposure to heat stress (37°C, 3 hours) and exposure to chronic oxidative stress (4 mM paraquat). We found that *sod-2* RNAi did not increase resistance to bacterial pathogens in *clk-1, clk-1;rde-1* and *clk-1* intestine-specific RNAi worms (**Figure 5A**). In contrast, ubiquitous or intestine-specific knockdown of *sod-2* significantly increased survival during heat stress (**Figure 5B**). RNAi-mediated knockdown of *sod-2* in the intestine of *clk-1* worms resulted in a similar decrease in resistance to oxidative stress to ubiquitous *sod-2* knockdown in *clk-1* animals (**Figure 5C**). Combined, this indicates that decreasing *sod-2* levels in the intestine is sufficient to affect resistance to heat and oxidative stress in the whole organism.

**Figure 5.**
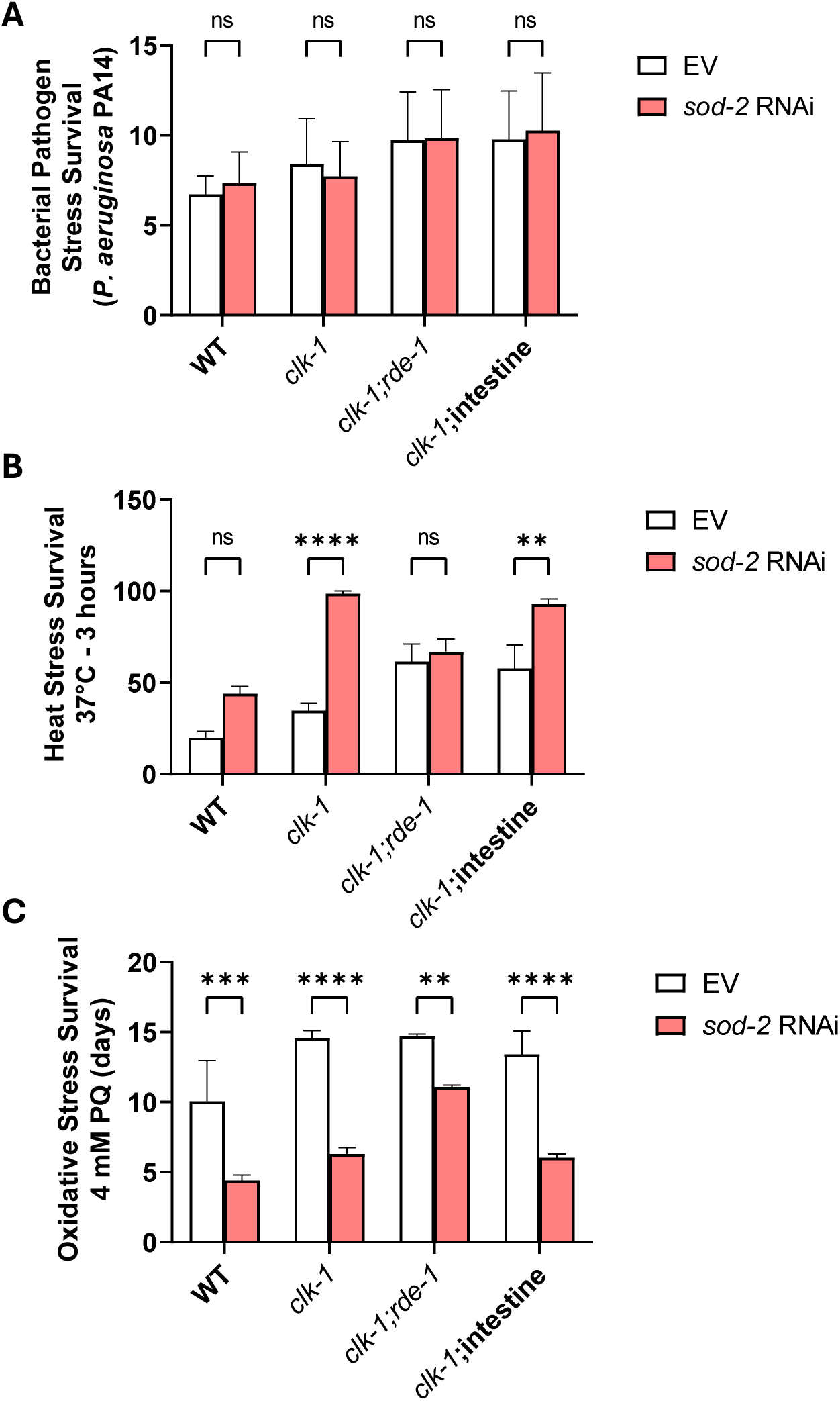
Intestine-specific knockdown of *sod-2* is sufficient to enhance resistance to heat stress. The effect of ubiquitous and intestine-specific *sod-2* knockdown on resistance to exogenous stressors was examined. (**A**) *sod-2* RNAi did not significantly increase resistance to bacterial pathogens (*P. aeruginosa* strain PA14) in WT, *clk-1, clk-1;rde-1* or *clk-1* worms with intestine-specific RNAi. (**B**) In contrast, knockdown of *sod-2* increased heat stress survival (37°C, 3 hours) in *clk-1* worms with ubiquitous RNAi or intestine-specific RNAi. (**C**) *sod-2* RNAi decreased resistance to oxidative stress (4 mM paraquat) to the same extent in *clk-1* animals with ubiquitous RNAi or intestine-specific RNAi. Three biological replicates were performed. Statistical significance was assessed using a log-rank test in panels A and C, and a two-way ANOVA with Šidák’s multiple comparisons test in panel B. *clk-1;* intestine = *clk-1;rde-1;kbIs7[nhx-2p::rde-1 + rol-6].* Error bars indicate SEM. **p<0.01, ****p<0.0001.

### Tissue-specific knockdown of *sod-2* alters physiologic rates

Deletion of *sod-2* decreases brood size, slows development, reduces thrashing rate and increases embryonic lethality [4]. To determine the extent to which these phenoyptes are caused by loss of *sod-2* in the intestine, we examined the effect of intestine-specific *sod-2* RNAi on physiologic rates. We found that knockdown of *sod-2* in *clk-1* worms markedly increased embryonic lethality (**Figure 6A**) while greatly reducing brood size (**Figure 6B**). In contrast, *sod-2* RNAi did not affect embryonic lethality or brood size in RNAi-inhibited *clk-1;rde-1* animals or *clk-1* worms in which RNAi is only active in the intestine (**Figure 6A,B**). Similarly, knockdown of *sod-2* greatly slowed the development of *clk-1* worms, but had little or no effect on the development of *clk-1;rde-1* or intestine-specific RNAi *clk-1* worms (**Figure 6C**). Finally, we found that *sod-2* RNAi did not affect thrashing rate in any of the strains tested (**Figure 6D**). Combined these results indicate that intestine-specific knockdown of *sod-2* increases lifespan in *clk-1* animals without altering physiologic rates.

**Figure 6.**
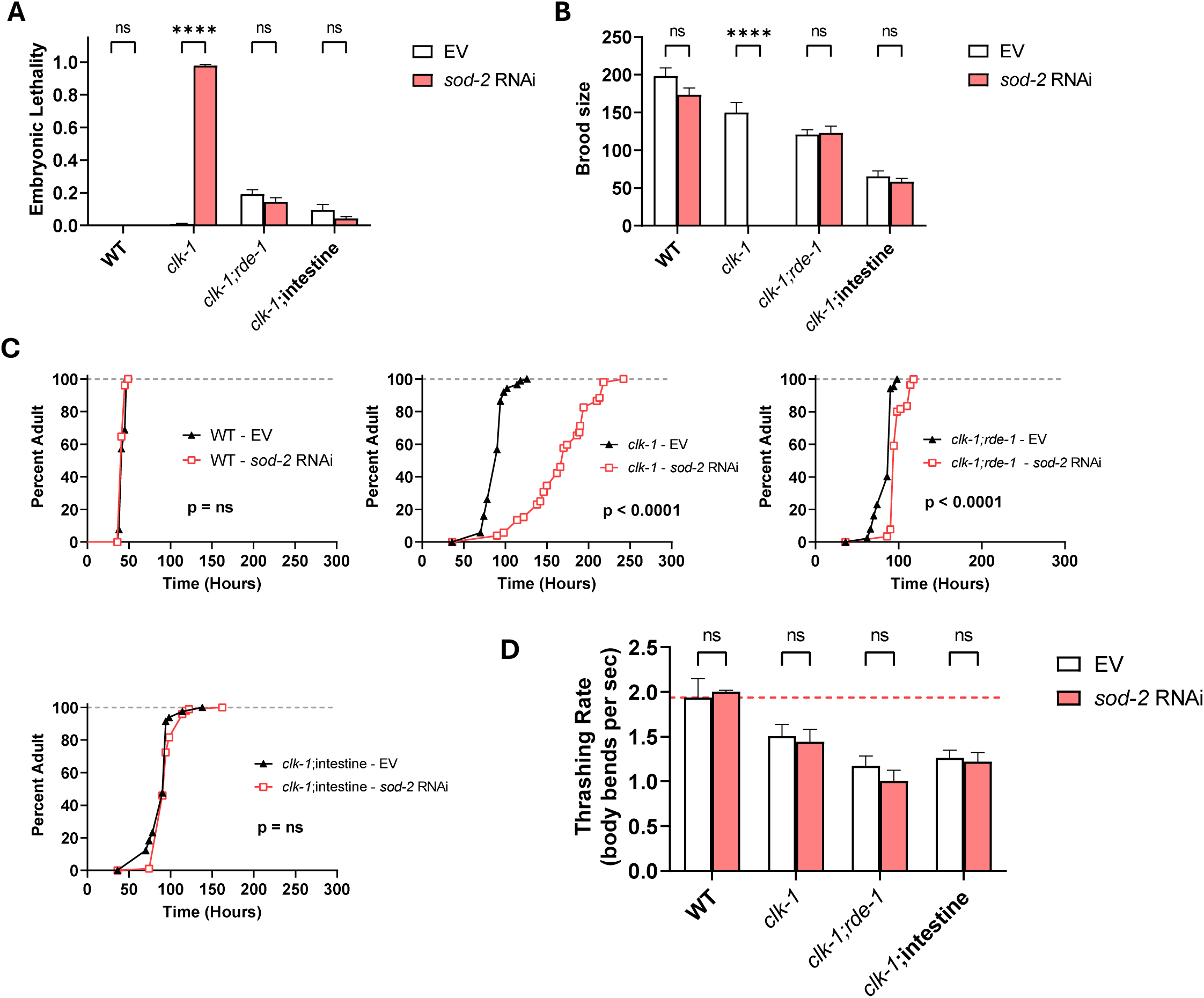
Intestine-specific knockdown of *sod-2* does not affect physiologic rates. The effect of ubiquitous and intestine-specific *sod-2* knockdown on physiologic rates was examined. (**A**) *sod-2* RNAi markedly increased embryonic lethality in *clk-1* worms with ubiquitous RNAi but had no effect in the intestine-specific RNAi strain. (**B**) Similarly, knocking down *sod-2* expression ubiquitously greatly reduced brood size in *clk-1* worms, while intestine-specific *sod-2* RNAi had no effect. (**C**) *sod-2* RNAi slowed the development of *clk-1* worms with ubiquitous RNAi, but had no effect when RNAi was limited to the intestine. (**D**) *sod-2* RNAi did not significantly affect the rate of thrashing in any of the strains tested. Three biological replicates were performed. Statistical significance was assessed using a two-way ANOVA with Šidák’s multiple comparisons test in panels A,B and D, and a log-rank test in panel C. *clk-1;* intestine = *clk-1;rde-1;kbIs7[nhx-2p::rde-1 + rol-6].* Error bars indicate SEM. ****p<0.0001.

## Discussion

Mildly increasing mitochondrial superoxide levels either indirectly through disruption of the mitochondrial SOD gene *sod-2* [4] or directly through treatment with the superoxide-generating compounds paraquat or juglone [6, 7, 40] results in increased lifespan. Interestingly, there are multiple other long-lived mutants that have elevated levels of ROS that contribute to their lifespan extension. The long-lived mitochondrial mutants, *isp-1* and *nuo-6,* both have elevated mitochondrial superoxide and show decreased lifespan when superoxide levels are decreased with the antioxidants N-acetyl cysteine (NAC) or vitamin C [5, 6]. Similarly, long-lived *daf-2* insulin/IGF-1 receptor mutants also have increased mitochondrial superoxide and reduced longevity in the presence of antioxidants NAC or butylated hydroxyanisole (BHA) [6, 8]. Most recently, it has been shown that long-lived germline ablation models have increased levels of ROS and that their lifespan is decreased by lowering ROS levels with the antioxidant vitamin C [9].

All of these long-lived genetic mutants provide the opportunity to dissect the mechanisms contributing to aging and lifespan extension. In order to target molecular investigations into the mechanisms involved, it is important to elucidate the tissue or tissues involved. In this work, we identified the tissues that are necessary or sufficient for a mild increase in mitochondrial superoxide to increase lifespan.

### Tissues necessary for lifespan extension

In order to determine which tissues are necessary for lifespan extension in *sod-2* mutants, we re-expressed wild-type *sod-2* in specific tissues using tissue-specific promoters. While ubiquitous expression of wild-type *sod-2* reduced the lifespan of *clk-1;sod-2* worms to the level of *clk-1* worms, expression of *sod-2* in germline, muscle, intestine or neurons did not decrease *clk-1;sod-2* longevity (**Table S1**). This indicates that there is no specific tissue among the tissues that we examined in which disruption of *sod-2* is necessary for lifespan extension. It is possible that *sod-2* disruption in a tissue that we did not examine is necessary to increase longevity. It is more likely that *sod-2* needs to be re-expressed in multiple tissues to prevent the lifespan-extending effects of global *sod-2* deletion.

While unable to rescue longevity, tissue-specific re-expression of *sod-2* was able to affect physiologic rates and stress resistance. Germline re-expression of *sod-2* reverted brood size, embryonic lethality and bacterial pathogen resistance towards control, indicating the *sod-2* deletion is acting in the germline to affect these phenotypes. Similarly, neuronal re-expression of *sod-2* reverted embryonic lethality, bacterial pathogen resistance and heat stress resistance caused by ubiquitous disruption of *sod-2* towards control.

In examining the insulin/IGF-1 signaling pathway, it was found that re-expression of *daf-2* in the neurons of *daf-2* mutants reduced lifespan to a similar extent as ubiquitous *daf-2* re-expression [23]. Re-expression of *daf-2* in the intestine resulted in a small decrease in *daf-2* lifespan while no effect was observed in muscle. Similarly, re-expression of *age-1* in the neurons or intestines of *age-1* mutants decreased lifespan towards wild-type [41]. The lifespan extension resulting from the disruption of the C-terminal binding protein CTBP-1 requires *ctbp-1* disruption in the neurons but not the hypodermis [42]. Neurons are also involved in the long lifespan that results from axenic dietary restriction, disruption of calcineurin and constitutive activation of AAK-2/AMPK. In the case of axenic dietary restriction, lifespan extension was decreased by knocking down *cbp-1,* which encodes the worm homolog of p300/CREB-binding protein, in neurons, or to a lesser extent, in germline or intestine, while no effect was observed in muscle, hypodermis or uterus [43]. In the case of calcineurin, expression of an activated form of the CREB-regulated transcriptional coactivator CRTC-1 in neurons, but not intestine, prevents lifespan extension resulting from disruption of calcineurin (*tax-6* RNAi) or constitutive activation of AAK-2 [44]. Finally, it has been shown that re-expression of the JNK homolog KBG-1 in neurons restores the decreased lifespan of *kgb-1* mutants to wild-type while expression in intestine, muscle or hypodermis has partial or no effect [45]. Combined, these previous studies highlight an important role for neurons in determining lifespan.

### Tissues sufficient for lifespan extension

To identify the tissues in which disruption of *sod-2* is sufficient to extend lifespan, we used tissue-specific RNAi. Using the approach, we found that knocking down *sod-2* in the intestine is sufficient to extend lifespan, while RNAi against *sod-2* in the neurons, muscle, germline or hypodermis did not increase longevity. This suggests that *sod-2* disruption in the intestine results in cell non-autonomous effects on the longevity of the whole organism. Intestine-specific *sod-2* RNAi was also sufficient to enhance resistance to heat stress and reduce oxidative stress resistance, but had no effect on bacterial pathogen resistance, embryonic lethality, brood size, development time or thrashing (**Table S2**), indicating that the effect of *sod-2* disruption on these latter phenotypes can be experimentally dissociated from its effect on lifespan.

In testing the sufficiency of DAF-16 to extend the lifespan of *daf-2;daf-16* mutants, re-expression of *daf-16* in the intestine increased lifespan while no effect was observed for neurons or muscle [24]. Similarly, expression of *daf-16* in the intestine of *mes-1;daf-16* double mutants resulted in a complete restoration of lifespan extension [24]. Tissue-specific knockdown of *daf-2* using RNAi in neurons, intestine and to a lesser extent hypodermis and germline was sufficient to increase lifespan, but not as much as ubiquitous *daf-2* RNAi [46]. Interestingly, neuronal knockdown of *daf-2* required intestinal *daf-16* to increase lifespan while intestinal knockdown of *daf-2* needed neuronal *daf-16* to extend longevity, suggesting that communication between neurons and the intestine mediate the lifespan extension caused by decreased insulin/IGF-1 signaling. It was also shown that intestine-specific degradation of *daf-2* using the auxin-degron system is sufficient to extend lifespan while degradation of *daf-2* in neurons, hypodermis germline had a much smaller effect, and no effect was observed in muscle [47].

In examining the tissue-specificity of other pathways, it was shown that overexpression of the heat shock factor HSF-1 in muscle, intestine or neurons all resulted in a mild increase in lifespan [48]. Similarly, expression of DAF-18 in the neurons, intestine or muscle of *daf-2;daf-18* worms was sufficient to increase lifespan [49]. For mild impairment of mitochondrial function, it was found that knocking down the cytochrome-c oxidase gene *cco-1* in intestine or neurons could increase lifespan while no effect was observed in muscle or hypodermis [20]. Interestingly, for the endoplasmic reticulum unfolded protein response, expression of a constitutively active version of the X-box binding protein gene *xbp-1* in the neurons or intestine is sufficient to increase lifespan, while ubiquitous expression or expression in muscle has no effect or decreases lifespan respectively [50]. Similar to the insulin/IGF-1 pathway, the unfolded protein response appears to extend lifespan through interaction between the neurons and intestine as disruption of specific lysosomal genes in the intestine inhibited the lifespan increase resulting from expression of constitutively active XBP-1 in neurons [51]. Finally, it was shown that tissue-specific knockdown of Mondo complex genes *mml-1* or *mxl-2* in neurons reduces the extended long lifespan of *daf-2* mutants and *glp-1* germline ablation mutants, while overexpression of *mml-1* in neurons is sufficient to increase the lifespan of wild-type worms [52]. Combined with our results, these studies demonstrate that both the intestine and neurons are key tissues for extending longevity and at least in some cases these two tissues appear to be working together.

Our work further suggests that levels of mitochondrial ROS, negatively controlled by SOD-2, can have beneficial effects in the intestine and detrimental effects in the neurons and germline of *C. elegans*. Consistently, rescuing SOD-2 in neurons and germline alleviated embryonic lethality caused by global *clk-1;sod-2* while extending lifespan further. Rescuing SOD-2 in the germline also improved the loss of fertility in *clk-1;sod-2* mutants. Meanwhile, rescuing SOD-2 in the intestine exacerbated embryonic lethality, impaired time to maturity, and slowed movement, whereas knockdown of *sod-2* in the intestine extended lifespan.

## Conclusions

Overall, this work examines the effect of tissue-specific re-expression of *sod-2* in animals lacking *sod-2*, and tissue-specific knockdown of *sod-2* in animals with wild-type *sod-2* expression. While tissue-specific re-expression of *sod-2* did not inhibit lifespan extension, it did revert stress resistance, fertility and embryonic survival towards control worms. Knockdown of *sod-2* in the intestine is sufficient to extend lifespan and alter resistance to stress but does not enhance longevity in any other tissue. Defining the conditions under which elevated mitochondrial superoxide increases lifespan will aid continued studies into the molecular mechanisms involved by targeting the tissues in which key molecular events are taking place.

## Acknowledgments

Some strains were provided by the CGC, which is funded by NIH Office of Research Infrastructure Programs (P30 OD010440). We would also like to acknowledge the *C. elegans* knockout consortium and the National Bioresource Project of Japan for providing strains used in this research.

## Author Contributions

Conceptualization: JVR. Methodology: TL, SZ, MS, SJT, JVR. Investigation: TL, SZ, MS, SJT, UA, AT, SKS, JVR. Analysis: TL, SZ, MS, SJT, JVR. Visualization: TL, SZ, MS, SJT, JVR. Validation: TL, SZ, MS, SJT, JVR. Writing – original draft: TL, JVR. Writing – review and editing: TL, SZ, MS, SJT, UA, AT, SKS, JVR. Supervision: JVR.

## Conflict of interest

The authors declare that no conflicts of interest exist.

## Materials & Correspondence

Correspondence and material requests should be addressed to Jeremy Van Raamsdonk.

## Funding

This work was supported by the Canadian Institutes of Health Research (CIHR; http://www.cihr-irsc.gc.ca/; JVR; Application 399148 and 416150) and the Natural Sciences and Engineering Research Council of Canada (NSERC; https://www.nserc-crsng.gc.ca/index_eng.asp; JVR; Application RGPIN-2019-04302). JVR is the recipient of a Senior Research Scholar career award from the Fonds de Recherche du Québec Santé (FRQS) and Parkinson Quebec. TL, AT, and SKS received studentship awards from the FRQS. SJT received a Vanier scholarship from CIHR. The funders had no role in study design, data collection and analysis, decision to publish, or preparation of the manuscript.

---

**Figure S1.**
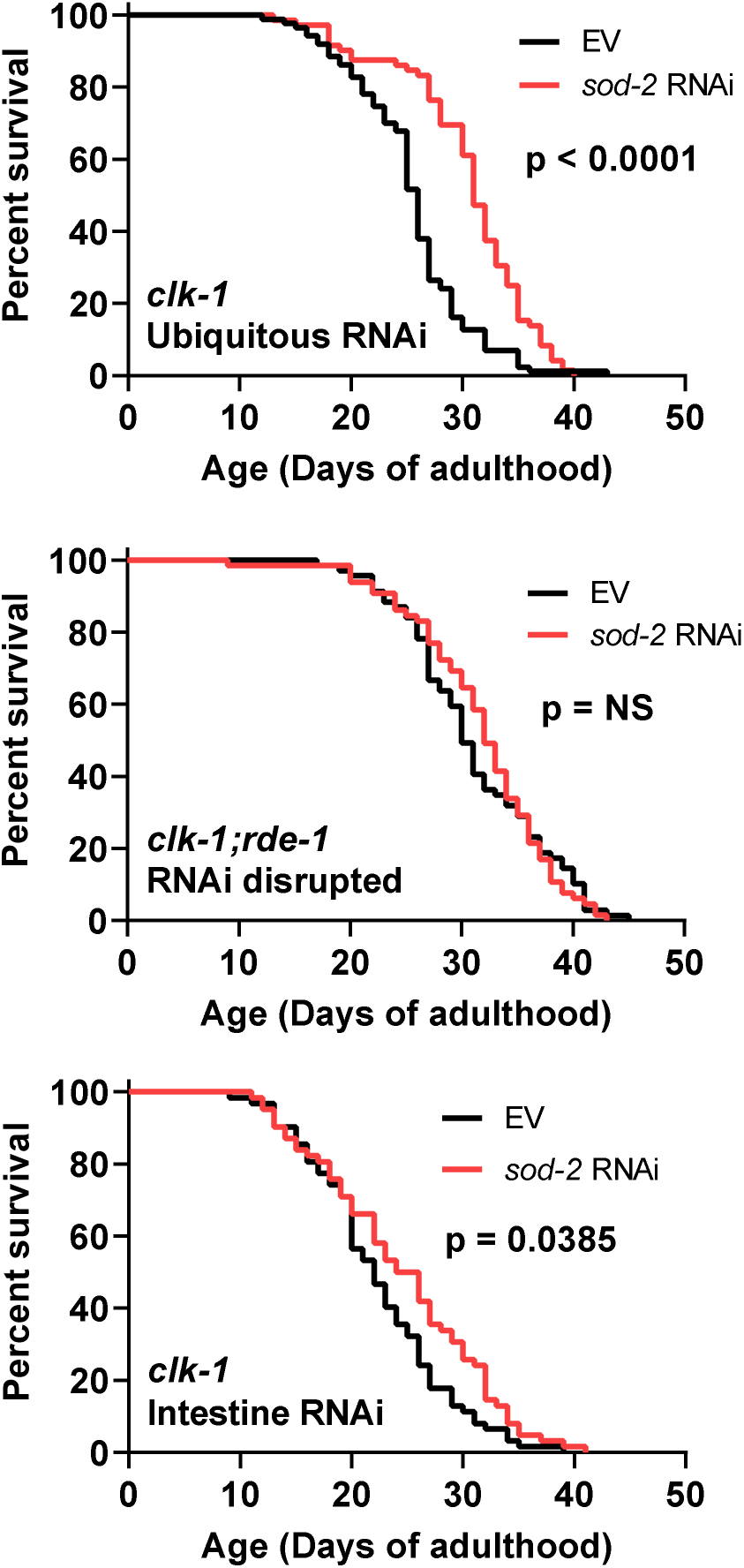
Mild elevation of mitochondrial superoxide in the intestine is sufficient to increase lifespan. To confirm the ability of intestinal *sod-2* knockdown to increase lifespan, we performed three additional replicates for *clk-1, clk-1;rde-1* and *clk-1* worms with intestine-specific RNAi. As in the original experiment *sod-2* RNAi significantly increased the lifespan of *clk-1* worms and *clk-1* worms in which RNAi is only active in the intestine but not *clk-1;rde-1* worms in which RNAi is inhibited. Three biological replicates were performed. Statistical significance was assessed using the log rank test. Intestine RNAi = *clk-1;rde-1;kbIs7[nhx-2p::rde-1 + rol-6]*.

**Table S1.** Summary of *sod-2* re-expression experiments. Arrows indicate the direction of significantly significant changes: ↑ indicates an increase, ↓ indicates a decrease, = indicates no effect. Green highlighting indicates that the phenotype was modulated towards the control indicating that *sod-2* re-expression is reverting the phenotype caused by *sod-2* deletion.

|  | Lifespan | Brood size | Embryonic lethality | Thrashing rate | Development time | Bacterial pathogen resistance | Heat stress resistance | Oxidative stress resistance |
| --- | --- | --- | --- | --- | --- | --- | --- | --- |
| Ubiquitous | ↓ | ↑ | ↓ | = | ↓ | ↓ | = | ↓ |
| Germline | ↑ | ↑ | ↓ | = | = | ↓ | = | = |
| Muscle | ↑ | = | ↑ | = | = | = | ↓ | ↑ |
| Intestine | = | = | ↑ | ↓ | ↑ | = | = | ↑ |
| Neurons | ↑ | = | ↓ | = | = | ↓ | ↓ | ↑ |

**Table S2.** Summary of intestine-specific *sod-2* knockdown experiments. Arrows indicate the direction of significantly significant changes: ↑ indicates an increase, ↓ indicates a decrease, = indicates no effect. *clk-1*; intestine = *clk-1;rde-1;kbIs7[nhx-2p::rde-1 + rol-6]*.

| Phenotype | Effect of *sod-2* knockdown |  |  |  |
| --- | --- | --- | --- | --- |
|  | WT | *clk-1* | *clk-1;rde-1* | *clk-1*;intestine |
| Lifespan | = | ↑ | = | ↑ |
| Bacterial pathogen resistance | = | = | = | = |
| Heat stress resistance | = | ↑ | = | ↑ |
| Oxidative stress resistance | ↓ | ↓ | ↓ | ↓ |
| Embryonic lethality | = | ↑ | = | = |
| Brood size | = | ↓ | = | = |
| Development time | = | ↑ | =/↑ | = |
| Thrashing rate | = | = | = | = |

## Notes

### Competing Interest Statement

The authors have declared no competing interest.

